# Metagenomic profiling of bacterial endosymbionts in wild mutant and permethrin-susceptible head lice shows an expanded microbiota in the resistant strain

**DOI:** 10.64898/2026.07.14.738380

**Authors:** Jalal Mohammadi, Hamzeh Alipour, Kourosh Azizi, Mohsen Kalantari, Mohammad Djaefar Moemenbellah-Fard

**Author notes:** Corresponding Authors: Mohammad Djaefar Moemenbellah-Fard; Research Center for Health Sciences, Institute of Health, Department of Biology and Control of Disease Vectors, School of Health, Shiraz University of Medical Sciences, Shiraz, Iran; Hamzeh Alipour. Mobile: +98917.

## Abstract

**Background:** Human head lice (*Pediculus humanus capitis* de Geer) are known to harbor diverse maternally inherited bacterial symbionts. These endosymbiotic bacteria may contribute to insecticide degradation, potentially helping lice withstand particular environmental pressures. Using next-generation sequencing (NGS), this study investigated the bacterial symbionts present in wild head-louse populations and characterized their phylogenetic relationships in mutant and permethrin-susceptible strains.

**Methods:** Head lice specimens were collected from 10 locations across Fars Province, Iran. Following DNA extraction, the samples were analyzed using polymerase chain reaction (PCR), and the resulting amplicons were sequenced to detect mutations in specimens from each location. Lice were then classified according to the presence or absence of mutations and subjected to NGS to characterize the symbiotic bacterial communities in mutant and putatively permethrin-susceptible strains.

**Results:** Mutant strains were detected at only three sampling stations. Bioinformatic analysis of the nucleic acid sequences revealed three exons and two introns, with an expected amplicon length of 582 bp. NGS analysis showed that *Candidatus* Riesia pediculicola (*Arsenophonus*), belonging to the phylum Proteobacteria, was the predominant bacterial genus. Actinobacteria and Firmicutes were the second- and third-most abundant phyla, respectively. Most of the remaining 25 bacterial taxa were associated with the mutant strain. Additionally, two previously unreported bacterial genera were deposited in GenBank.

**Conclusions:** The distinct distribution of *Arsenophonus* species between susceptible and mutant head-lice strains—along with the greater abundance of *Escherichia*, *Shigella*, *Lawsonella*, and *Megamonas* in mutant strains—highlights the need for advanced metagenomic analyses to determine how these endosymbionts may help their hosts withstand specific environmental disturbances.

## Background

Metazoan animals, including head lice, coexist with a wide variety of both useful and harmful microbes (1). It is mostly believed that intricate symbiotic partnerships have opened new ecological niches conducive to the incredible diversification of insects (2, 3). Pediculosis capitis as a condition due to the head lice, *Pediculus humanus capitis* de Geer, 1767 (Anoplura: Pediculidae), has been one of the most frequently irritating human infestations worldwide since primordial past. Though this obligatory infestation appears rather asymptomatic to humans, it causes intense pruritus and is intolerable to many people in the high-income countries of the world (4).

Various therapeutic protocols have been formulated to combat this infestation, including physical, herbal and chemical interventions (5, 6). Chemical procedure is the main treatment line globally, among which applies lotions bearing insecticides. The widespread use of over-the-counter (OTC) pediculicides, lack of proper treatment, or incomplete and unprofessional intervention have recently led to lice resistance against some of them. The control of this infestation has thus become burdensome. In recent years, lice resistance to treatment has increased globally, and various factors have contributed to this, the most significant of which has been gene exchange between lice (7, 8). The short life cycle of louse, its relatively small genome (9), the transfer of genetically modified lice species from infested areas to other parts of the world and the use of more or less permissible pediculicides have led to the unimpeded access in genetic transmission between lice, and ultimately these genetic changes have generated lice resistant to treatment (5, 10).

Though pediculosis capitis does not categorically cause infectious disease in *Homo sapiens*, and the fact that head lice cannot still be ascertained to have the vector capacity of transmitting any pathogens (11), the potential presence of prokaryotes and their possible role in lice resistance to insecticides have recently been implicated. This phenomenon has recently been demonstrated in obligate blood feeding bed bugs and other insects (12–17). Head lice can ingest pathogens such as *Bartonella quintana* (18), *Borrelia recurrentis*, and *Rickettsia prowazekii* (the causative agents of trench fever, relapsing fever, and epidemic typhus, respectively), and then discharge them onto the body surface (19). These pathogens may spread diseases such as bacillary angiomatosis, chronic lymphadenopathy, and endocarditis to humans (20). However, head lice have a stronger immune response than body lice, and lead to eliminating ingested pathogens (21).

Recent studies, however, have shown that head lice may embody a few pathogens of humans, including *Acinetobacter* sp., *Bartonella quintana*, *Coxiella burnetii*, *Borrelia recurrentis*, *Borrelia theileri* and *Yersinia pestis* (22, 23). The DNAs of new species of *Anaplasma* and *Ehrlichia* have also been extracted from head lice (24). Given that head lice could be implicated in possible harborage of pathogens for humans, and are a cause of widespread human infestation worldwide, identification and control of these pathogens seems indispensable to contain their versatility. Nevertheless, the mechanisms for resistance of pathogens to the immune response of lice remain unknown (22).

Since mammalian blood is a rich source of proteins, iron and salts, but almost devoid of lipids, carbohydrates and vitamins, obligatory blood feeding insects such as lice depend on symbiotic bacteria for supplementation of B vitamins (25, 26). The only limiting nutrient consistently linked with the evolution of obligate symbiosis was B vitamins (2). The endosymbiotic bacteria residing in the midgut of host louse commonly supply these nutrients to its feeding niche. Detecting microbial agents in lice is usually done by DNA extraction.

One of the DNA extraction and sequencing methods is the next-generation sequencing (NGS). This method has been widely used due to its cheapness and speed. NGS platforms perform extensive parallel sequencing, in which millions of DNA fragments are harmonically sequenced. The NGS method is a technique that conducts 16S rRNA sequencing on the bacterial genome (27, 28). It is the standard method for studying hybrid microbial populations.

In microbiology, NGS identifies the genome of pathogens. It means that instead of using conventional methods such as morphology, staining, and using metabolic properties to identify pathogens, NGS helps to identify and examine the relationship of pathogens with each other and track the source of infection and drug susceptibility of pathogens by defining the genome of pathogens (29). Recently, NGS has been used to identify bacteria that exist in lice. This method identifies the bacteria in lice by extracting and analyzing the DNA sequence (30).

The global prevalence rate of head louse infestation is about 19%, as reported in a recent meta-analysis-based systematic review (31). As outlined above, identifying head lice infection with various microbial agents is of particular significance to appreciate their likely role in deviating control measures. However, it is unclear to evaluate whether head lice resistance to treatment affects them harboring different endosymbionts. Therefore, this study aimed to examine and identify the infection with microbiota in mutant and permethrin-susceptible head lice in Fars province, Iran, using the NGS method.

## Materials and Methods

### Sampling

All methods were performed in accordance with the relevant guidelines and regulations under the institutional code of ethics (IR.SUMS.SCHEANUT.REC.1400.011). This research did not directly involve human participants or the use of live animals in experiments. Sampling of head lice were conducted by the regional health officers in the presence of parents and school nurses with a fine-toothed plastic detection comb (PDC) that was utilized to isolate them. The delivered head lice samples were washed thrice in double distilled water, and placed in 75% alcohol. They were then kept in -20°C freezer at School of Health, Shiraz University of Medical Sciences (SUMS). Lice gender and maturity were identified under a stereomicroscope.

### DNA Extraction and Processing

The DNA extraction from the whole body of lice after homogenization and addition of protease buffers was performed using SinaClon Kit according to the manufacturer’s protocol and kept at a temperature of -20°C. The samples were sent to Pishgam Tehran Company for sequencing. As a result, the sequences were analyzed. After the mutants were identified, the next stage was implemented.

The number of pools (35 head lice each) was two. The first pool of extracted DNA from natural head lice was mutant and the second one was sensitive to permethrin. DNA extraction was conducted on head louse populations from Marvdasht (29°87′87″N, 52°82′06″E) and Kazerun (29°62′71″N, 51°65′18″E) counties of Fars province. Exactly 20µl of each of the original frozen DNA samples from Marvdasht and Kazerun counties were sent to PARS Gene Fanavaran company, Tehran and South Korea (MacroGen). Symbiotic bacteria were detected in mutant and susceptible samples using NGS method, and metagenomic analyses were done using Illumina MiSeq device.

### Input Data

Overall, five template samples were received for 16S rRNA sequencing from tick models. The details about the samples are in Table 1 below. DNA extraction was dispatched to MacroGen, South Korea, for sequencing, then using 341F forward primer and 850R reverse primer (primer sequences were CCTACGGGNGGCWGCAG, and GACTACHVGGGTATCTAATCC, respectively) the DNA was amplified and subsequently sequenced with MiSeq device.

**Table 1.**
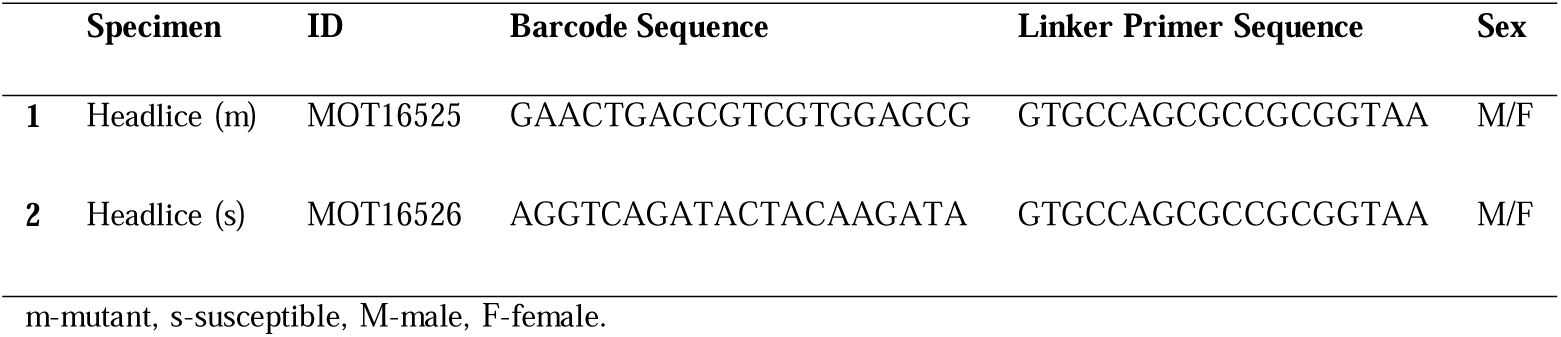
The metadata frame of head louse samples.

### Metadata Frame

The sequencer was set to sequence the PCR product of 300 base pairs from both ends.

### Joining and Denoising Reads

The DADA2 software was used to denoise and join the forward and reverse reads. Trimming lengths for forward and reverse reads are crucial in this step. Since the used primers were 341F and 850R, the expected length of amplicons was 850-341=509, so with paired end read length equal to 300, we have 2 x 300 - 509 = 91 bases overlap without trimming. It was recommended to leave 20+ natural amplicon variance bases for overlap and trim the rest for best results. Checking with FastQc results, we set the trim length for forward reads at 300 (no trimming) and for reverse reads at 250 (trimming 50 bases from the end). Since the length of target region (509) was much higher than the read length (300), the risk of having adapter sequence at the end of forward reads was negligible. The resulting operational taxonomic units (OTUs) were then annotated with SILVA database assigning each sequence to the nearest species that shared at least 90% of the sequence FastQc along with MultiQc was used to check the raw Fastq files for quality (Figure 1).

**Figure 1.**
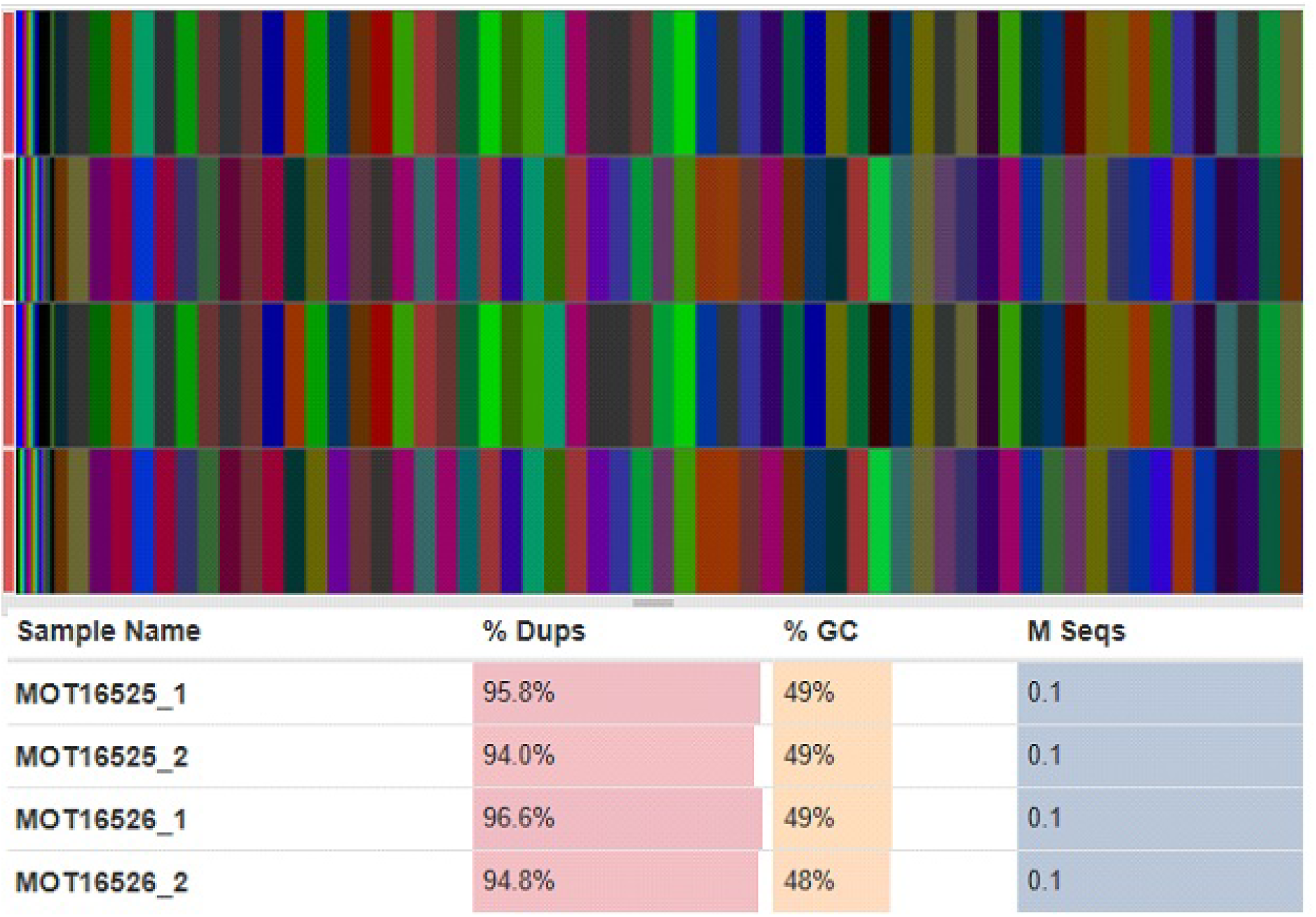
The raw Fastq files for quality control check. 1-Forward, and 2-reverse primers for the mutant (the last two digits 25) and susceptible (the last two digits 26) head lice, Dups-Duplicate reads, GC-guanine/ cytosine, M Seqs-milionth sequences.

### Phylogeny

The OTU sequences of mutant head lice from Marvdasht copied to MEGA 0.6 software and aligned by ClustalW. The phylogenetic tree was then drawn. The statistical method with Maximum Likelihood (ML). Bootstrap replication at 1000 duplicates was applied for the phylogeny tree. This tree was depicted based on the identified head louse specimens that were registered in NCBI and samples included: *Chitinophaga* sp. (OL654754.1), *Terrimonas* sp. (HM124372.1), *Novosphingobium clariflavum* (MN629069.1), *Novosphingobium resinovorum* (MN629068.1), *Klebsiella variicola* (OM432577.1), *Enterobacter hormaechei* (OM424283.1), *Phnomibacter ginsenosidimutans* (NR_173663), *Novosphingobium* sp. (MT240612.1) and *Entomoneis* sp. (MF997419.1).

## Results

The present study was performed on head lice collected in Dec. 2021 from Marvdasht and Kazerun counties in Fars province of south Iran. Samples were collected according to the code of ethics IR.SUMS.SCHEAUT.REC-1400.011 approved at SUMS. After sequences were ready, an analysis of bioinformatics sequences revealed the samples of Marvdasht County had three *kdr* gene mutations in sodium channels at V875L-Q876P-S879V positions. These were thus identified as mutant, and samples from Kazerun County without any mutations were regarded as sensitive to permethrin. DNA extracted by the NGS method entered the sequencing stage. This process was performed by Illumina MiSeq device and the output of this method was to obtain features, OTUs, and their amounts.

The samples of our research using the NGS program were related to the extracted DNA of permethrin-sensitive and -mutant head lice. In total 29,173 OUTs were outlined in both head lice strains, a smaller (46.8%) portion of which were from the mutant strain, while the larger (53.2%) portion belonged to the susceptible strain. The highest percentage of bacteria obtained were from Proteobacteria, Actinobacteria, Firmicutes and Bacteroidetes taxa. The preliminary data from both susceptible and permethrin-resistant head lice exhibited in a decreasing order of frequency that the phyla of Firmicutes (46%), Bacteroidetes (27%), Proteobacteria (15%), and Actinobacteria (12%) were the most abundant bacterial taxa.

Some 13 bacterial species belonging to the two genera of *Bacteroides* and *Lactobacillus* were demonstrated from the mutant category. The most prevalent (98.2%) bacterial genus was *Candidatus* Riesia pediculicola (represented here by *Arsenophonus*). This highly abundant taxon in both sensitive and mutant categories was *Arsenophonus*, ambiguous. Only a minor percentage (≈ 3.7% and ≈ 0.2% in mutant and susceptible categories, respectively) of bacteria in the two categories were different (Figure 2). There were disparities in the number of Proteobacteria, which was 2335 more in susceptible than mutant species and 282 *Escherichia*-*Shigella*, which was present in mutant non-susceptible species (Table 2). The total number of bacteria counted was 29,173 of which 13,654 (46.8%) bacteria lied in the mutant category that included 96.34% of OTUs (Figure 2) and were mostly related to the first phylum (*i.e.* Proteobacteria), and 3.66 % of bacteria were related to the other taxa that were present in the first (mutant) and not in the second susceptible category. In the second category, 15,519 (53.2%) bacteria were counted. It has been found that 99.81 % (Figure 2) were again related to the first taxon as above and only 0.19 % were related to bacteria present in the second (susceptible) and not in the first mutant category.

**Figure 2.**
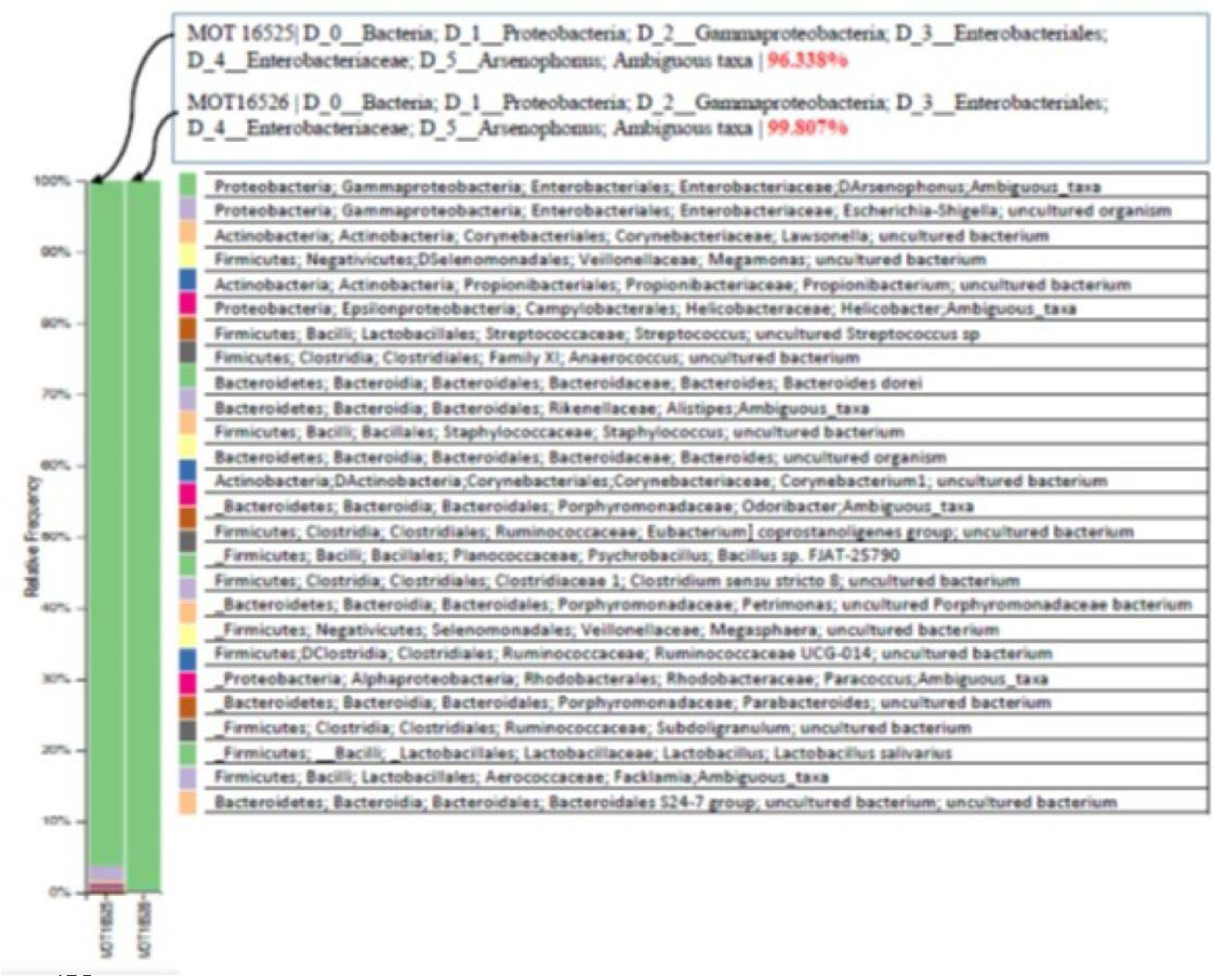
Relative frequency (%) of bacterial communities at the class level in mutant (MOT16525) and susceptible (MOT16526) head lice samples.

**Table 2.**
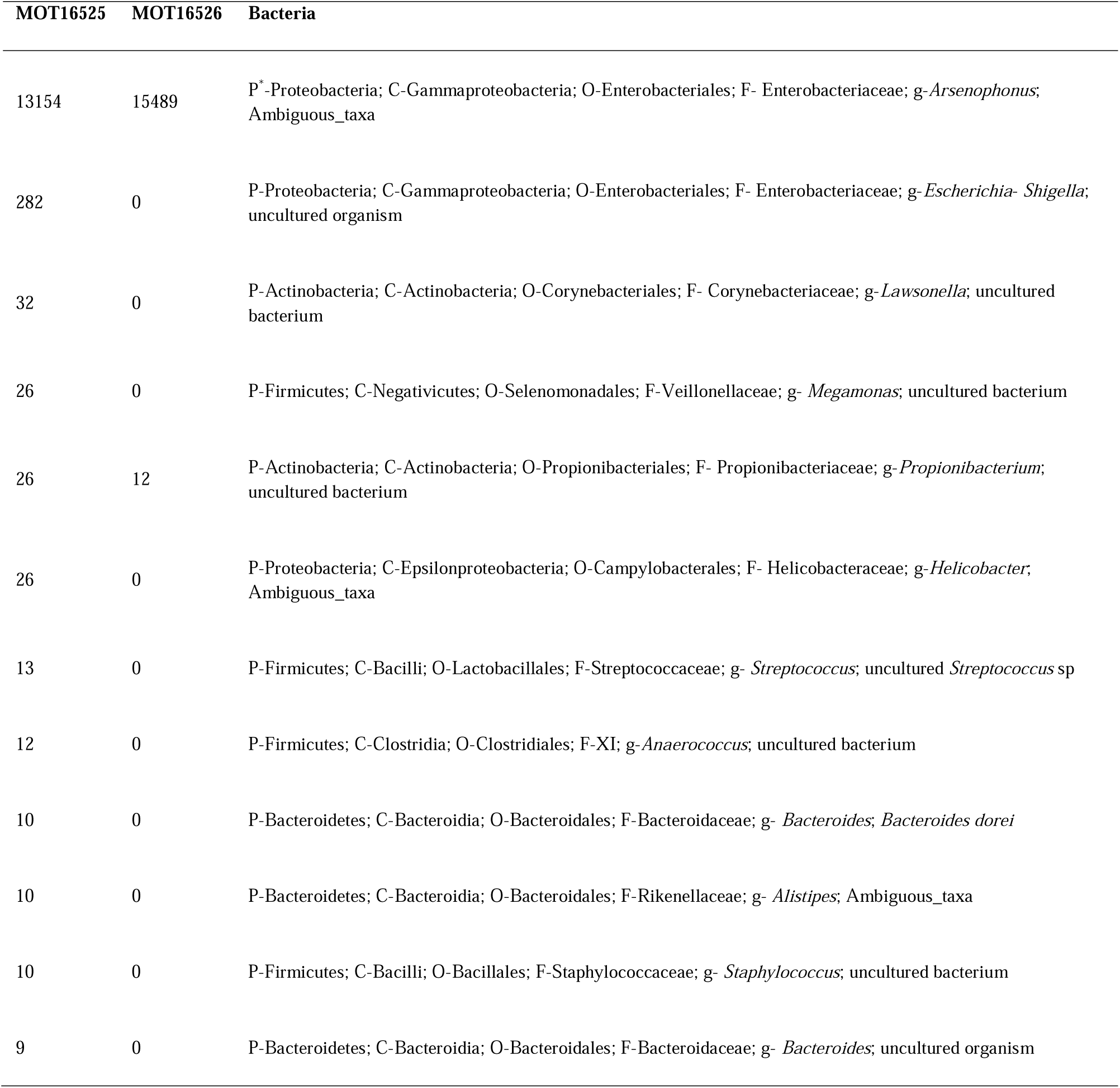
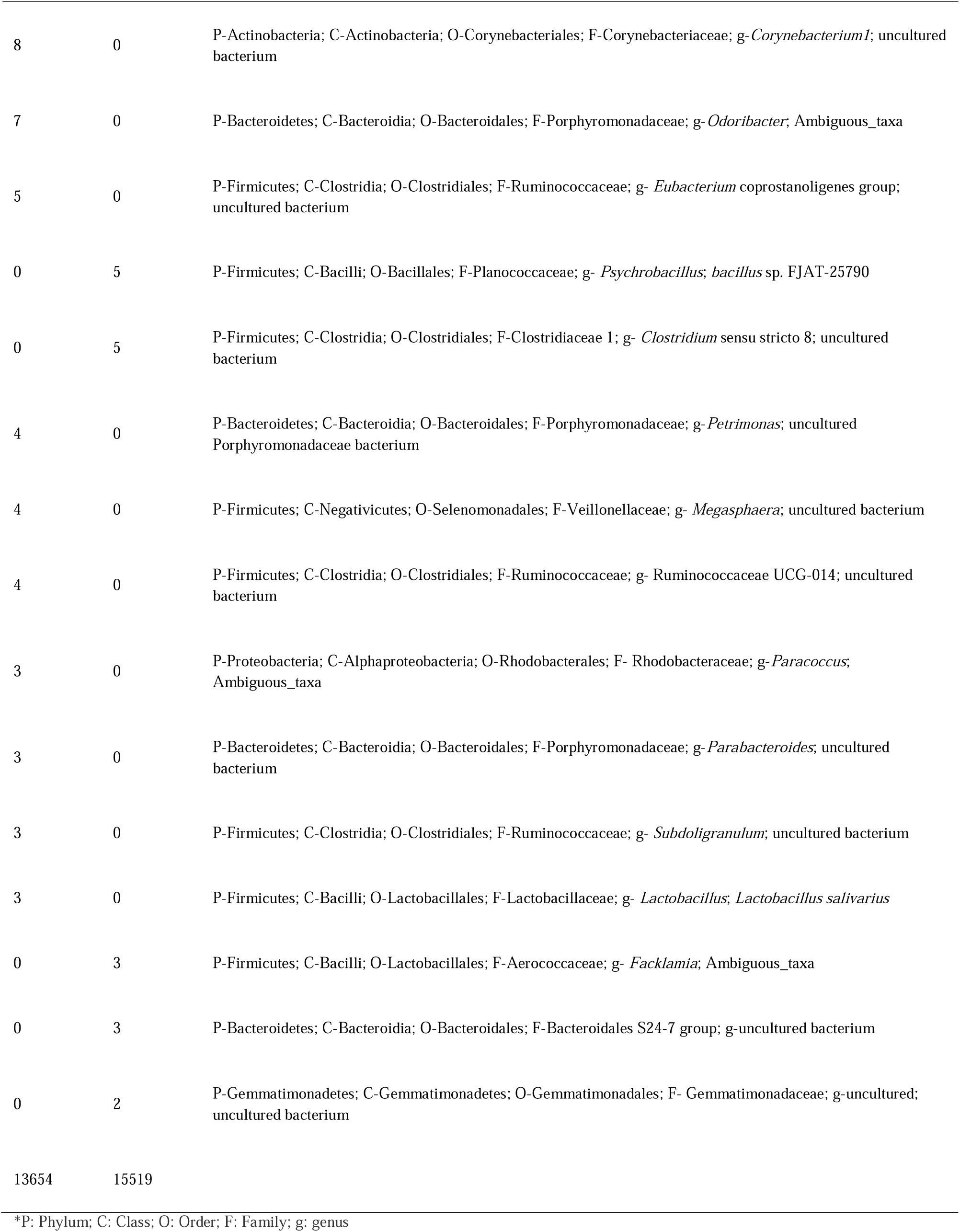
Abundances of various OTUs in the mutant (1^st^ left column) and susceptible (2^nd^ left column) categories of head lice.

The difference between the number of bacteria in categories 1 (mutant) and 2 (susceptible) was discernible in front of each classification of bacteria. For instance, there were 26 numbers in the mutant sample and 12 numbers in the sensitive sample from *Propionibacterium* taxon (Table 2).

Both samples were similar in 96.34% of bacterial OTUs belonging to the mutant category including the phylum of Proteobacteria. Except the two common genera of *Arsenophonus* and *Propionibacterium* from the mutant and susceptible lice, all other bacterial genera were only present in the mutant forms. New strains of bacterial endosymbionts have also been identified in the NGS analysis that have been recorded unprecedentedly (Table 3).

**Table 3.**
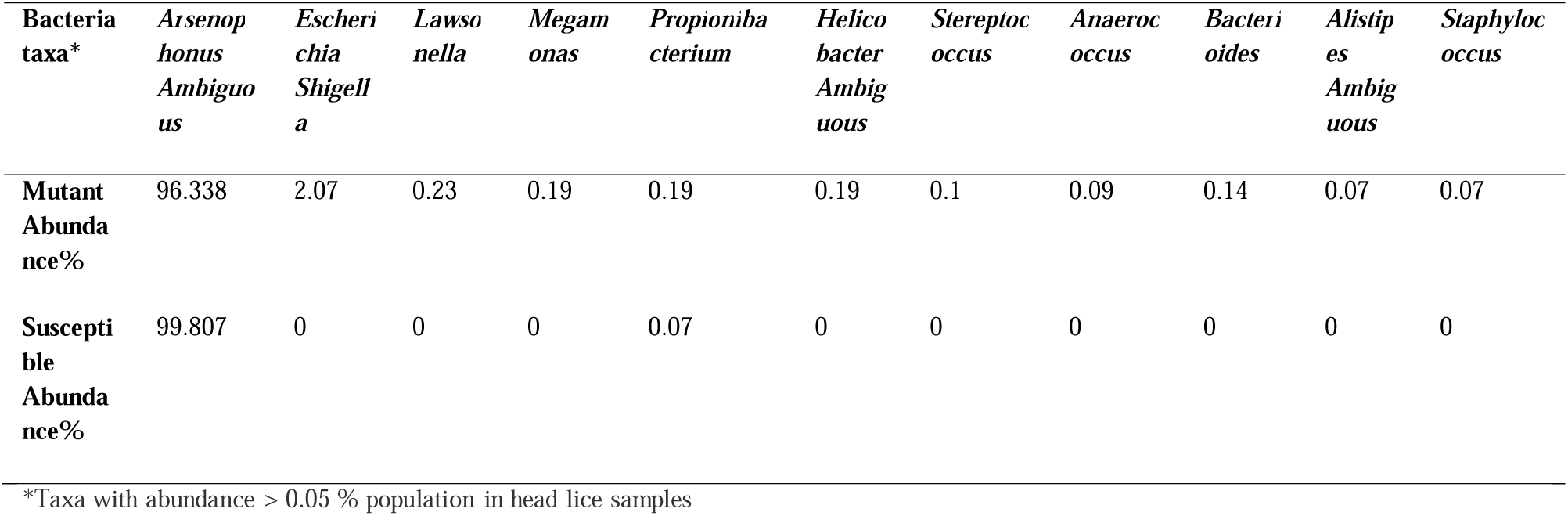
Comparative abundance of bacterial genera within head lice samples.

### Phylogeny

The 16S rRNA sequence analysis between the mutant head lice bacterial species found in this study and other bacteria obtained from the GenBank was shown in Figure 3. From the phylogenetic tree, OTU1 clustered into the *Entomoneis* sp. (MF997419.1). Clades. OTU2 and OTU5 grouped with *Novosphingobium clariflavum* (MN629069.1), *Novosphingobium resinovorum* (MN629068.1) and *Novosphingobium* sp. (MT240612.1). OTU3 and OUT6 clustered together with *Terrimonas* sp. (HM124372.1) and *Phnomibacter ginsenosidimutans* (NR_173663); in addition, OUT4 clustered with *Klebsiella variicola* (OM432577.1) and *Enterobacter hormaechei* (OM424283.1) (Figure 3).

**Figure 3.**
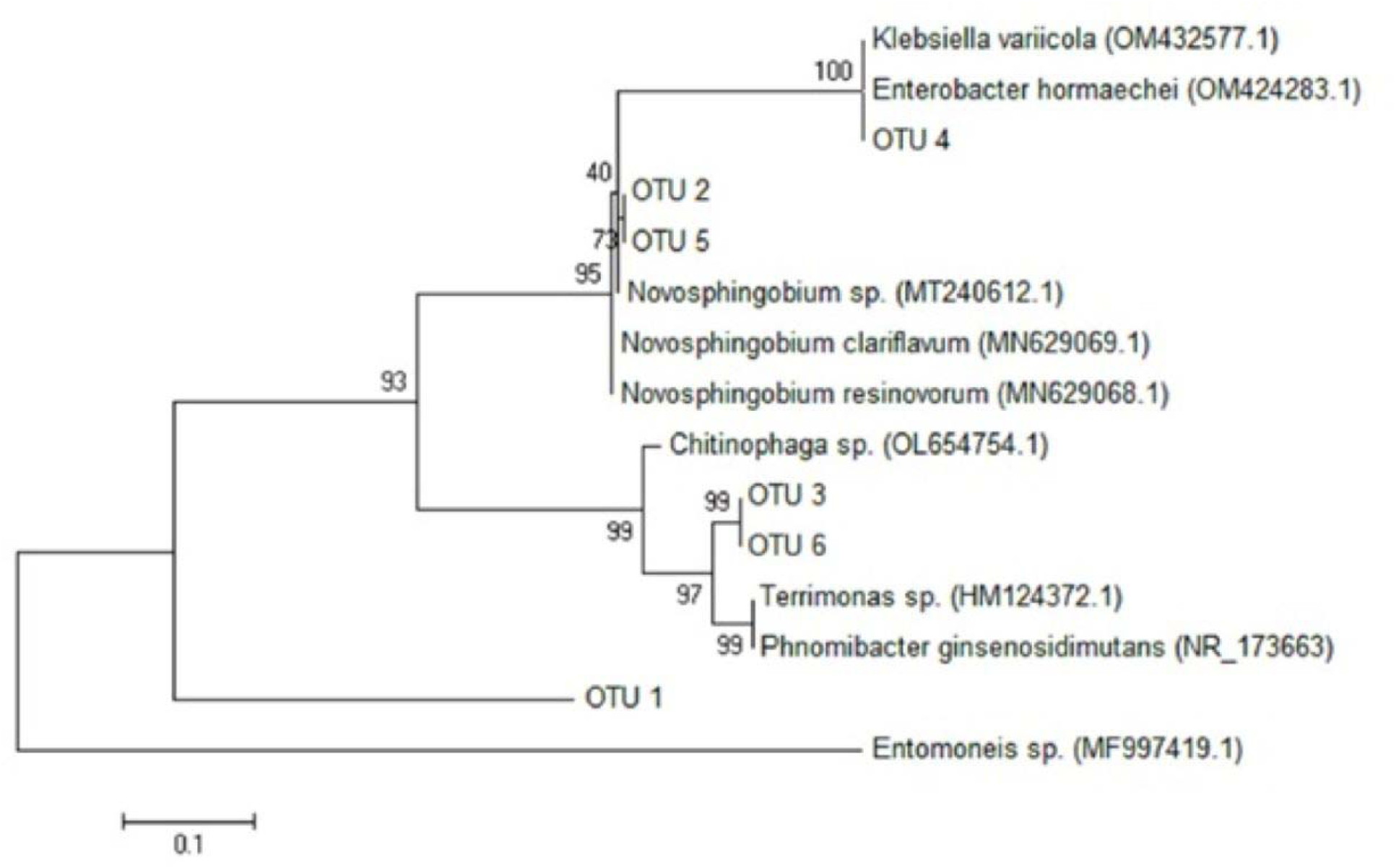
Phylogenetic tree from 16S rRNA sequence isolated from mutant head lice. In the present study, OTUs were compared with other bacterial strains in GenBank. The phylogenetic tree was drawn by the Maximum Likelihood (ML) method. Phylogeny tree was indicated by the 1000 bootstrap replicates. GenBank accessions numbers for reference sequences are shown at the end of any naming.

## Discussion

The preceding results revealed that mutant head lice harbored not only a wide diversity of opportunistic bacteria which were absent from susceptible ones, but that there were also nine newly-identified GenBank registered bacteria whose presence in the mutant head lice of Marvdasht county provided challengeable scenarios to be contemplated. The preliminary outcome from both susceptible and permethrin-resistant head lice showed that the phyla of Firmicutes (#12), Bacteroidetes (#7), Proteobacteria (#4), and Actinobacteria (#3) were the most frequent bacterial taxa in a descending order of preponderance.

These were also the most prevalent symbiotic bacterial taxa in the intestinal tract of all larval black soldier flies (BSF), *Hermetia illucens* (32) and the forensically-important pupal stages of the blowfly species, *Chrysomya megacephala* (33). The promotive effects of these diverse phyla on the growth of BSF were analyzed. Likewise, the presence of 16 different microorganisms belonging to the two different phyla of Firmicutes and Proteobacteria was somehow reminiscent of those found in the bloodsucking sand fly insect, *Phlebotomus spp.*, vectors of leishmaniasis in the Old World (12, 34). It indicated that the host habitat interplay could shape the variable diversity of housefly (35) and sand fly gut microbiome (12).

The most abundant endosymbiotic bacterial genus was *Candidatus* Riesia pediculicola (*Arsenophonus*), since this endosymbiont is ubiquitously present across populations of lice. This is the most commonly widespread intracellular symbiont in many insect orders including Hymenoptera, Diptera, Hemiptera and Phthiraptera (36). The symbiont *Arsenophonus* usually lies inside midgut specialized host cells to locate symbiotic bacteria, called bacteriocytes (37, 38). These gut-linked bacteriocytes-housed *Arsenophonus* clade could move from stomach discs to ovarian ampulla to be transferable (via maternal or transovarial transmission) by developing eggs to the next generation of lice (38). An analysis of more than a hundred *Arsenophonus* sequences from 54 host taxa exposed its roughly monophyletic clade, illustrated by unique molecular synapomorphies (39).

The higher prevalence of merely this genus in susceptible lice and its intimate association with *Escherichia-Shigella, Lawsonella,* and *Megamonas* bacterial genera in mutant lice was noticeable. The increased preponderance of microbiota (*Bartonella quintana*, *Acinetobacter* spp.) in susceptible than resistant lice has also indirectly been demonstrated in permethrin-resistant head lice from Georgia compared to those of susceptible body lice from Russia (10, 40).

Most microbiota, including bacteria, coexist with arthropods for nutrition (*e.g.* the essential B-group vitamins (37), growth, and survival, and act as a reservoir for them. Lice are host-specific arthropods that carry numerous endosymbionts with them. A limited blood diet of head louse can provide essential nutrients for the growth and development of bacterial endosymbionts, so finding a relatively high number of pathogens, especially *Arsenophonus, Rhodococcus* and *Pediculicola* species in lice is normal (41–43). However, it is common to see other types of microsymbionts whose relevance to lice has hitherto remained unknown.

The prolific heritable endosymbiont *Arsenophonus* that was highly widespread in both head lice strains of this study could belong to different clades. Both primary/obligate (P) and secondary/facultative (S) endobionts depend on their relevant genome sizes (*i.e.* a diminished genome of few hundred kb compared to a large one of a few Mb reflect P-or S-symbiotic roles, respectively) linked with their specific hosts (37). P-symbionts are usually transferable vertically, while S-symbionts move laterally between host vectors. It is conceivable that some *Arsenophonus* species of lice could adopt P-vs. S-endobiont role to preserve or levitate the fitness and survival of itself and its host species.

There is some speculation that a small genome size confers long evolutionary history with the host insect lineage (37), and the subsequent deletion of many unessential genes causing genomic erosion (44, 45). Convincingly, a larger genome size provides for a broader metabolic potential producing a wider range of essential nutrients, such as thiamine (B1), riboflavin (B2), niacin (B3), pantothenic acid (B5), pyridoxine (B6), biotin (B7) and folate (B9) (46).

Furthermore, evidence from an *Arsenophonus*-aphid system indicates that this endobiont could itself receive further protection from a lysogenic bacteriophage that encodes a range of homologous chemicals (36), which could be capable of enduring resistance against certain environmental stresses such as insecticide use. This bacteriophage may readily move between quite distant symbionts horizontally, and be a swift and strong mechanism of immediate adaptation for endobionts and their host insects. According to a recent report, the divergence between the *Arsenophonus* (endosymbiont of *Lipoptena cervi*, the deer louse flies) and the symbiont, *Riesia* clades, dates back to 13-25 million years ago (18).

In addition, 16S rDNA sequences revealed a strict co-evolution between the endosymbionts of the boar louse, *Haematopinus apri*, compared with those of the hog louse, *Haematopinus suis* (25). Based on their endosymbiont sister clades congruity, as the domestic pig originated from the wild boar more than 8000 years ago following its domestication, a similar scenario could be suggested for the divergence of human body lice from head lice after the advent of clothing and civilization (22, 25). Others reported, however, a minimum of 107,000 years for the origin of clothing (47). A recent evidence corroborates the fact that the endosymbionts phylogeny generally mirrors the human host louse phylogeny with a clear co-phylogenetic congruence over time, reflecting a strict transovarial transmission and a host louse-symbiotic bacteria co-speciation following the evolutionary process of the human louse (18).

The *Shigella*-*Escherichia* symbiotic bacteria (the second in frequency after *Arsenophonus*) that are also small gram-negative non-motile rod in the tribe Escherichieae, family Enterobacteriaceae, were only present in the mutant strains of head lice in this study. A likely speculation is that these bacillary symbionts could confer gene mutations leading to the permethrin resistance in head lice directly through enzyme activities or indirectly through bacteriophages.

Eventually, it is an inescapable topic that despite the unequivocal tempo of research in the field of controlling communicable diseases in recent years, the challenge of ameliorating vector-borne zoonotic, and specifically anthroponotic, maladies remains a troublesome task to resolve (48).

## Conclusions

Investigating the microbiota of obligate ectoparasites may provide novel approaches for addressing insecticide resistance. Expanding our understanding of their endobionts could clarify differences in bacterial composition and reveal potential contributions to insecticide resistance in wild-caught susceptible and mutant head louse strains. The bacterial taxa identified in human head lice from southern Iran were similar to those previously reported in neighboring countries, including Turkey and Azerbaijan.

## DECLARATIONS

### Ethics approval

This study attained its appropriate ethical approval under the following code: IR.SUMS.SCHEANUT.REC.1400.011, from SUMS.

### Consent for publication

All co-authors have their full consent for this MS to be published.

### Data availability statement

All data are concisely reported in the main MS transparently. Further information on data could be made available upon request from the corresponding authors. The datasets generated and/or analyzed during the current study are available in the repository “[https://mega.nz/file/TrJAVTxC#QGwz6BAOesIdOvmFLxI9rAQFYGEMP8oqXsTBvL35U8g]”.

### Authors contributions statement

J.M., H.A., and M.D.M-F. designed the protocol. H.A. and J.M. carried out the experimentation. M.D.M-F., M.K., and K.A. consulted on the software handling and progress of work. H.A., J.M., and M.D.M-F. analyzed the data and wrote the pre-draft. M.K. and K.A. helped with figures. All authors read and reviewed the final MS.

## Acknowledgements

This manuscript was the final part of a Ph.D. thesis by Mr. Jalal Mohammadi (Grant number: 22579). All authors are grateful to the support provided by the vice-chancellor for research and technology at SUMS, Shiraz, Iran. This research would not have been conceived without the active participation of other research staff at Shiraz School of Health and the publisher’s respected reviewers and editor-in-chief of the relevant journal, to all of whom we are indebted.

